# Proteomics and Ex Vivo Plaque Culture Identify the Insulin-Like Growth Factor Axis as a Regulator of Carotid Plaque Stability

**DOI:** 10.1101/2025.10.04.680463

**Authors:** Sara M. Jørgensen, Karen C. Yang-Jensen, Karin Yeung, Timothy A. Resch, Jonas Eiberg, Michael J. Davies, Lasse G. Lorentzen

## Abstract

**Objective:** Rupture of carotid atherosclerotic plaques leading to cerebral embolization, is a significant cause of stroke. We previously analyzed 21 plaques by mass spectrometry and reported that the proteomes of morphologically unstable (rupture-prone) and stable plaques are different. This dataset extends and includes non-atherosclerotic (thyroid) arteries to allow comparison with control tissue, and to investigate plaque stability using *ex vivo* plaques cultured.

**Methods:** Plaques (n=76) and non-atherosclerotic superior thyroid artery segments (n=8) were retrieved from carotid endarterectomies. Additionally, 22 plaques were cultured *ex vivo* for 22 days to examine the role of insulin-like growth factor-1 (IGF-1) signalling. Proteins were analyzed by liquid chromatography-mass spectrometry.

**Results:** Mass spectrometric proteome analysis identified three protein clusters associated with morphologically unstable (type A) and stable (type B) plaques, as well as non-atherosclerotic arteries. 2,876 proteins were differentially abundant in plaques compared to non-atherosclerotic arteries. 1,415 proteins were differentially abundant between plaque types A and B. Proteins linked to IGF transport and binding, particularly IGF-binding proteins, were more abundant in plaques compared to non-atherosclerotic arteries, and in type B compared to type A plaques. IGF-1, IGF-2 and the IGF-1 receptor were more abundant in type B plaques, whereas the IGF-2 receptor was more abundant in type A. IGF-1 treatment of *ex vivo* plaques decreased matrix metalloprotein 9 and increased collagen type XXI, consistent with a increased plaque stability.

**Conclusions:** Proteomic analyses of atherosclerotic plaques, and *ex vivo* plaques cultured with IGF-1, reveals the IGF axis as a potential regulator of human atherosclerotic plaque stability.

**CLINICAL RELEVANCE:** The protein composition of unstable carotid artery plaques differs from that of stable ones, which may explain their varying tendency to rupture. Components of the insulin-like growth factor (IGF) axis are more abundant in atherosclerotic plaques than in healthy tissue, and are more abundant in stable compared to unstable plaque morphology, suggesting a protective role of these proteins against plaque rupture. To investigate this, we treated plaques *ex vivo* with IGF-1. This data indicates that manipulating the IGF axis may promote plaque stability, with potential clinical relevance in prevention of plaque rupture.

**ARTICLE HIGHLIGHTS:** *Type of Research:* Human study

*Key Findings:* Proteomic analysis of 76 carotid atherosclerotic plaques and 8 superior thyroid artery controls identified the insulin-growth factor (IGF) axis as potential regulators of plaque stability. *Ex vivo* culture and IGF-1 treatment of 11 symptomatic carotid plaques induced proteome changes, compared to 11 controls, consistent with features of plaque stabilization.

*Take home Message:* The insulin-like growth factor axis is a potential regulator of carotid atherosclerotic plaque stability, as revealed by proteomic analysis and *ex vivo* plaque culture.

## INTRODUCTION

Thrombosis and embolization caused by plaque rupture in the carotid artery are an important cause of ischemic stroke. Unstable plaques are more rupture-prone and are characterized by a large lipid-rich necrotic core, a high level of inflammation, intra-plaque bleeding, a thin fibrous cap of extracellular matrix (ECM) due to diminished synthesis or enhanced degradation, few vascular smooth muscle cells (SMCs), and less calcification compared to stable plaques [1]. In a recent study, we analyzed 21 carotid artery plaques by mass spectrometry-based proteomics [2]. Comparison of the abundance of protein species indicated that plaques with macroscopic stable appearance, judged by the surgeon, were enriched in ECM proteins, whereas plaques with an unstable or intermediate appearance were enriched in proteins involved in inflammation and protein degradation.

Here we report an extended proteomic analysis of 76 carotid artery plaques from symptomatic patients and non-atherosclerotic ‘control’ artery samples from the superior thyroid artery, collected as paired controls from 8 patients. This approach enables comparative proteomic analyses between ‘healthy’ tissue and atherosclerotic plaques, as well as across different plaque subtypes. The aim of this study was to examine potential pathways associated with plaque stability, using both direct analyses and plaques cultured *ex vivo*.

## METHODS

### Ethics

This study was approved by The Danish National Committee on Health Research Ethics (journal number H-20002776) and conducted according to the Helsinki declaration. For the initial 76 patients (non-cultured plaques and paired control artery), patient data and biological materials were collected with patients’ written, verbal, and informed consent. In the case of the additional 22 plaques that were cultured, biological materials were collected as fully anonymised waste material that did not require ethical approval under Danish legislation, as per the decision of the Independent Regional Scientific Ethics Committee of the Capital Region of Denmark (case number: 23050906).

### Peroperative carotid endarterectomy tissue samples

Carotid plaques (n=76) from consecutive patients, recently diagnosed with cerebral embolic ischaemia (stroke, transient ischaemic attack) or retinal ischaemia (complete vision loss, amaurosis fugax) were collected and processed as previously described [2]. Briefly, plaques were removed *in toto* during eversion carotid endarterectomy (CEA) under locoregional anaesthesia. As part of the CEA procedure of the last 8 included patients, the thyroid artery was surgically dissected, ligated and excised. The superior thyroid artery is commonly used as non-atherosclerotic control tissue as reported in previously literature [3]. The carotid plaques and thyroid arteries were subsequently washed and stored at −80L in a preservative buffer. Plaques were classified immediately after surgery as *soft*, *hard*, or *mixed* by the operating surgeon, as previously described [2, 4]. Briefly, classification was based on the overall consistency of the plaque and macroscopic morphological features such as calcification, fibrosis, disintegrated content, intraplaque haemorrhage and ulceration. Macroscopic classification was performed prior to, and independently of, the proteomic analyses.

### *Ex vivo* plaque culture

Carotid artery plaques from an additional 22 patients were collected as described above. The plaque *in toto* was cut into cross-sectional rings (∼2-3 mm) and cultured in stable isotope-labelled amino acid (SILAC) medium for 22 days under standard cell culture conditions (see Supplementary Methods). SILAC media contains heavy isotope labelled versions of the ami-no acids lysine (Lys) and arginine (Arg). These heavy isotope-labelled amino acids will be incorporated into proteins during synthesis in the *ex vivo* cultured plaques, which induces a mass shift detectable by mass spectrometry. This permits monitoring of newly synthesized proteins in the *ex vivo* cultured plaques and the discrimination of proteins synthesized during *ex vivo* culture from unlabelled, light-isotope proteins present within the plaque prior to excision and culture. Half of the plaques (n = 11) were supplemented with 100 ng/ml IGF-1 with this added to the culture medium and renewed every third day.

### Proteomic analysis

Protein extraction, clean-up and digestion was carried out as described in previous work [2], except that clean-up and digestion were carried out using a robotic platform. Briefly, the protein component of the artery samples (either freshly stored or cultured *ex vivo*) was extracted and solubilized using the Sample Preparation by Easy Extraction and Digestion (SPEED) protocol [5], then neutralized. Proteins were reduced, alkylated, and captured on magnetic beads in the presence of acetonitrile, followed by washing with acetonitrile/ethanol and on-bead digestion with Lys-C and trypsin. The resulting peptides were separated from the magnetic beads, acidified and stored at −80 °C until analysis. Peptide samples were separated and analyzed using a Dionex Ultimate RSLCnano system equipped with an Aurora C18 column (15 cm×75 μm, 1.6 μm particle size, IonOpticks) coupled to a Bruker timsTOF Pro mass spectrometer. Peptides were eluted using a binary gradient (2-35% mobile phase B in 17.5 min followed by washing and equilibration of the column; mobile phase A: 0.1% v/v formic acid in MS-grade water, mobile phase B: 0.1% v/v formic acid in acetonitrile) with data acquired using data-independent acquisition (DIA) in parallel accumulation–serial fragmentation (PASEF) mode (DIA-PASEF) [6]. The workflow is graphically summarised in **Fig 1** and described in detail in the Supplementary Methods.

**Figure 1.**
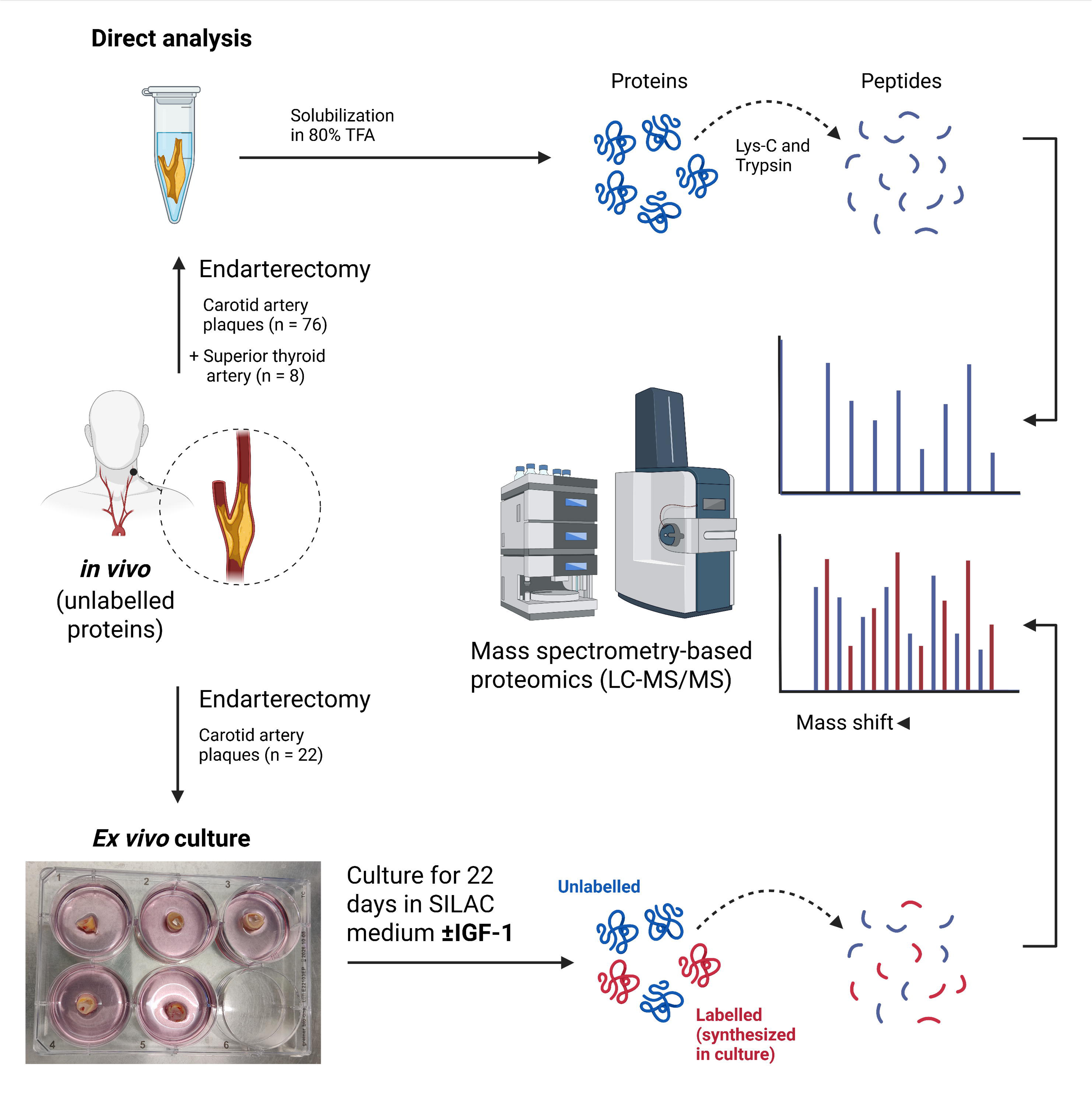
Overview of experimental workflow from sample collection to liquid chromatography-tandem mass spectrometry (LC-MS/MS) analysis. **(Top)** Plaques were obtained from CEA, and in the case of 8 patients, a section of the superior thyroid artery was also excised. Proteins were extracted, cleaned up, digested, and analysed by LC-MS/MS as described in Methods. **(Bottom)** In a parallel workflow, plaques were collected for *ex vivo* culture with and without IGF-1 supplementation. Plaques were cultured in SILAC medium for 22 days, resulting in labelling of newly synthesized proteins with isotopically labelled amino acids (Lys and Arg). This introduces mass shifts in proteins and peptides detectable by mass spectrometry, allowing for discrimination of proteins synthesized during *ex vivo* culture from (un-labelled) proteins present within the plaque prior to excision and culture. Proteins were extracted, cleaned up, digested and analysed by LC-MS/MS as for direct analysis above (described in Methods). This figure was created with Biorender.com.

### Data analysis and statistics

DIA-PASEF data were processed using the DIA-NN software [7, 8] using a library constructed from DDA-PASEF data obtained from analysis of off-line fractionated samples (see Supplementary Methods). Further data analysis and visualisation were performed in R. Summarisation of precursor signal to protein levels, and statistical inference with linear modelling was carried out using the MSqRob [9] workflow, with Benjamini–Hochberg adjusted *p*-values <0.05 considered significant [10, 11]. The Monte Carlo Reference-based Consensus Clustering (M3C) package [12] was used to cluster the samples using the k-means algorithm and assess the clustering patterns based on the 20% most variable proteins. Optimal cluster number was determined based on maximal Relative Cluster Stability Index (RCSI), an empirical *p*-value less than 0.05 (comparing stability scores to null distributions generated by Monte-Carlo simulation) and a minimal entropy calculated by the algorithm. Multidimensional scaling (MDS) was used to visualize the similarity between samples based on global protein expression profiles. MDS projects high-dimensional expression data into a two-dimensional space, such that the distance between points reflects the leading sources of variation between samples. The analysis was performed on the 500 most variable proteins using the plotMDS function from the *limma* R package. For differences between tissue types, Reactome pathway analysis was performed by enrichment analysis with log2 fold changes of protein intensity (differential abundance) or over-representation analysis with natural log odds ratios of precursor counts (differential detection) for ranking and scoring. Differences between plaque types (A and B) were only examined by enrichment analysis using log2 fold changes for ranking and scoring. Scripts to reproduce analyses and visualizations are provided in the Supplementary Data.

### Data availability

In accordance with Danish legislation relating to the General Data Protection Regulation (GDPR), raw mass spectrometry data can only be shared under a data processing agreement between the data controller and processor. Such data sharing agreements are available and can be completed by contacting the corresponding authors.

## RESULTS

Atherosclerotic carotid plaques (n=76) from 76 patients and 8 non-atherosclerotic sections from the superior thyroid artery (**Table I**) were analyzed by proteomics using the protocol outlined in **Fig 1**, resulting in the identification of 7,343 proteins using a 1% false-discovery rate (FDR) cut-off. After filtering out single peptide identifications and proteins with high missingness (>90%) across all 84 samples, 5,731 proteins were quantified. Unsupervised consensus clustering was employed to obtain unbiased sample partitioning of the proteomics data. The relative cluster stability index reached its maximum at K = 3, indicating optimal partitioning into three clusters. This partitioning was observed to be more plausible (*p* < 0.05) compared to the null hypothesis (“No clusters”). No other partitioning (number of clusters) of the samples was significantly more plausible than the null hypothesis (**Fig 2A**).

**Figure 2.**
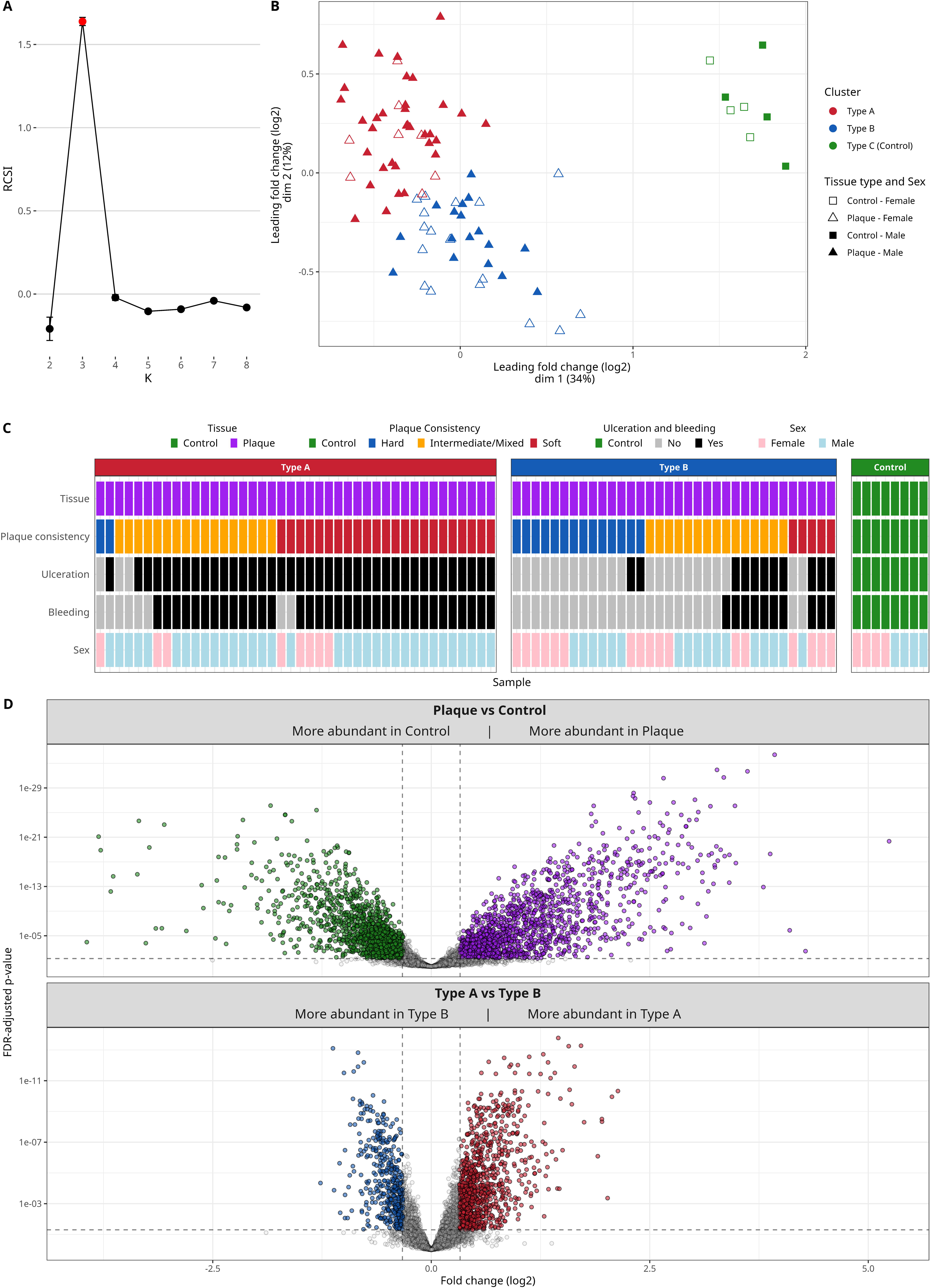
Classification of samples based on proteomic profiles as well as clinical parameters and comparison of protein abundances between identified clusters. (**A**) Relative cluster stability index (RCSI) values for clusters number (K) 2-8 obtained by Monte Carlo reference-based consensus clustering. Cluster numbers with significant Monte-Carlo *p*-value < 0.05 are represented by red dots, while non-significant cluster numbers are coloured black. (**B**) MDS plot of distances between protein abundance profiles for the control artery and plaque proteomes. Each sample is plotted on a two-dimensional scatterplot of the first two components of a MDS analysis of the top 500 most abundant proteins, with each shape representing an individual control artery (square) or plaque (triangle). Shapes are coloured based on their consen-sus, with open/closed symbols indicating the biological sex of the donor (open = female, closed = male). (**C**) Tile plot overview of all samples, including tissue type, consensus cluster, clinical parameters (plaque consistency, ulceration, and bleeding), and the biological sex of the donor. (**D**) Volcano plots showing the comparison of protein abundances between plaques and control tissue (top), and plaque clusters A and B (bottom). Each dot represents a protein with a log2 fold change between the samples compared (x axis) plotted against the FDR-adjusted *p*-value (reversed log10; y axis). Gray dots indicate non-significant changes (FDR-adjusted *p*-value > 0.05 with a cut-off set at a log2 fold change of ±0.33, which corresponds to a fold change of ±1.25).

Multidimensional scaling (MDS was used to assess and visualize overall similarities in protein abundances between samples and demonstrated a clear segregation of the clusters, consistent with a healthy control tissue group (type C) and two types of plaque (types A and B; **Fig 2B**). Plaques described as soft or intermediate/mixed based on the surgeon’s visual assessment, were predominantly associated with type A, whereas plaques described as hard or intermediate/mixed were associated with type B (**Fig 2C**). Using a Chi-squared test, we found a significant (*p* 0.009) association between patient biological sex and the consensus clusters, with a positive association detected between plaques from females and type B, while plaques from males were associated with type A.

Using robust linear modelling, we examined differences in protein abundance (based on intensity) and detection (based on precursor count) across the clusters (**Fig 2D**). Using a fold change cut-off of ± 1.25, 2,876 proteins exhibited significant differential abundance between control and plaque tissue (1,514 more abundant in plaques; 1,362 more abundant in controls). An additional 80 proteins were sparsely or not detected in controls but consistently detected in plaques. Comparison of the two plaque types indicated 1,415 differentially abundant proteins, with 986 more abundant in type A compared to type B, and 429 more abundant in type B compared to type A plaques.

### Proteins related to the regulation of IGF and IGFBPs are positively associated with plaques, and particularly type B (stable) plaques

To identify pathways and processes associated with plaques compared to control tissue, and between plaque types, Reactome pathway analyses were performed (**Fig 3**). Pathways linked to immune responses (including ‘toll-like receptor cascades’, ‘complement cascade’, and ‘neutrophil degranulation’), extracellular matrix organization, and regulation of IGF and IGFBPs were positively associated with plaque when compared to control tissue (**Fig 3**). Conversely, mitochondrial processes (‘aerobic respiration’ and ‘respiratory electron transport’) were negatively associated with plaques.

**Figure 3.**
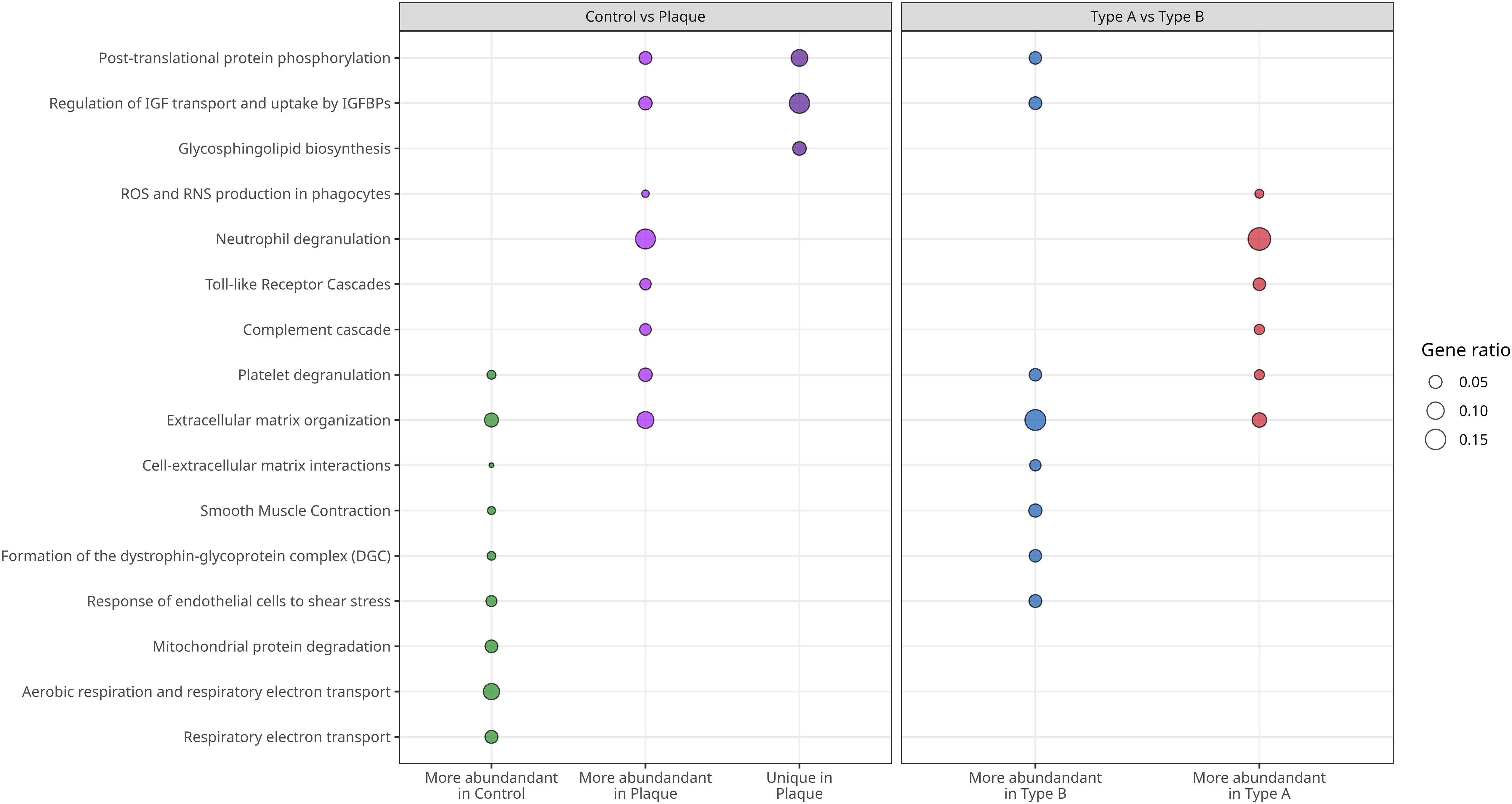
Reactome pathway analysis. Dotplot pathway enrichment map showing the most significantly overrepresented Reactome pathways (q < 0.05) when considering all differentially abundant/detected proteins. Left panel show results from comparing control tissue and plaque samples, with enrichments for proteins more abundant in control (green), more abundant in plaque (purple), and proteins only detected in plaques (dark purple) shown separately. Right panel show results from comparing type A and B plaques, with enrichments for proteins more abundant in plaque type B (blue) and plaque type A (red) shown separately. Sizes of dots are proportional to the percentage of proteins among differentially abundant/detected proteins belonging to the indicated pathway.

Comparisons between the plaque types indicated that pathways associated with immune responses were positively associated with type A plaques compared to type B. On the other hand, pathways related to respiratory processes, extracellular matrix organization, smooth muscle contraction, and regulation of IGF by IGFBPs were positively associated with type B (stable) plaques.

Whilst the association between inflammation with type A (unstable) plaques is unsurprising, fewer data are available on the IGF system [13]. As pathway(s) involving regulation of IGF by IGFBPs were positively associated with plaque tissue compared to healthy tissue, based on both differential abundance and detection, and differed between plaque types, these changes were examined in greater detail. On comparing plaque to control tissue, insulin growth factor-2 (IGF-2) and the binding proteins, IGFBP-1, IGFBP-2, IGFBP-3, IGFBP-5, IGFBP-6, IGFBP-7, and IGFBP acid-labile subunit (IGFALS) were significantly more abundant and/or detected more frequently. The IGF-1 receptor (IGF-1R) was less abundant and detected less in plaques, whereas the IGF-2 receptor (IGF-2R) was more abundant (**Fig 4A**). Comparison between the two plaque types, showed that IGF-1, IGF-2, IGFBP-1 and IGFBP-7 were more abundant and/or detected in type B compared to type A plaques. No differences in abundance or detection were found for other IGFBPs, though IGFALS was more abundant and more detected in type A plaques. For the receptors, IGF-1R was more detected in type B, whereas IGF-2R was more abundant in type A plaques (**Fig 4B**).

**Figure 4.**
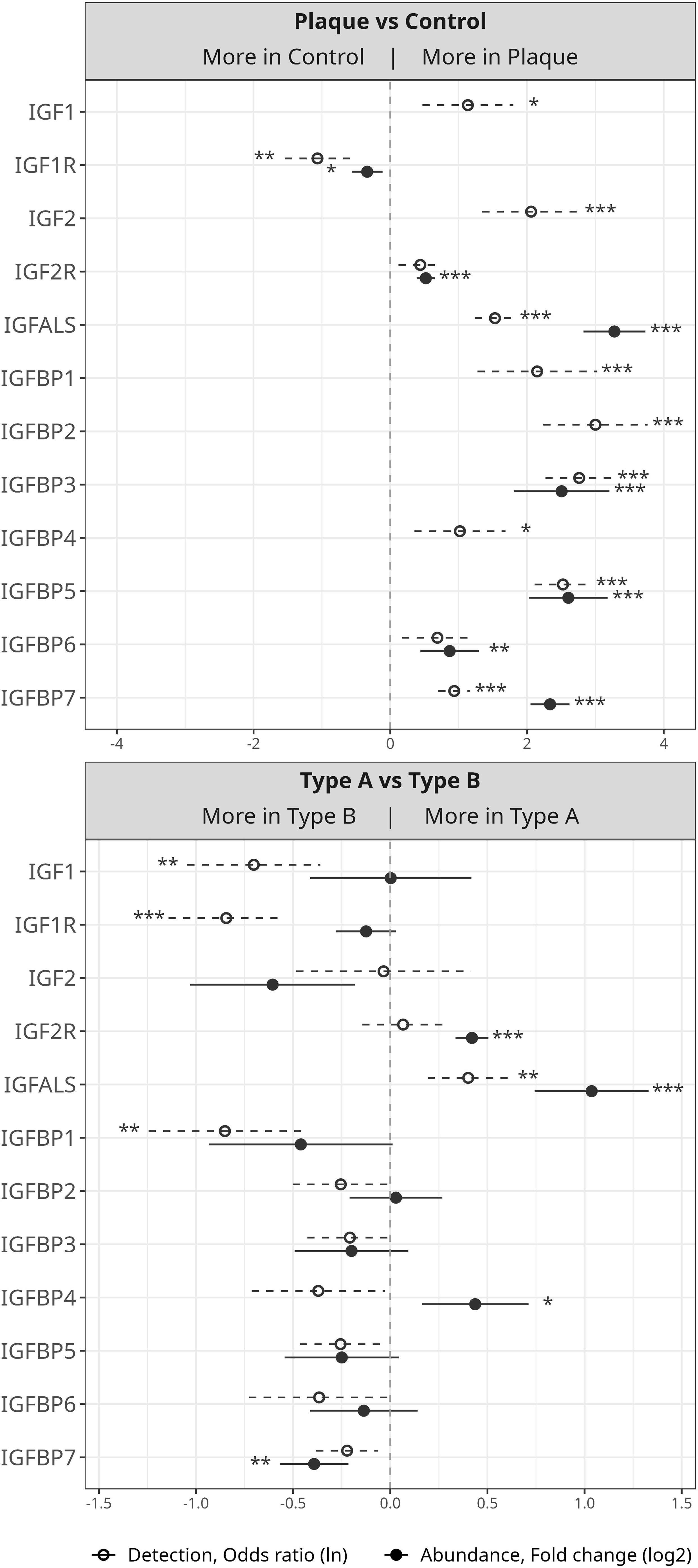
Abundance and detection of selected proteins of the ‘regulation of IGF transport and uptake by IGFBPs’ pathway. Forest plots displaying the detection (precursor count, natural log of odds ratio; orange) and abundance (intensity, log2 fold change; blue) and detection of selected proteins from the ‘regulation of IGF transport and uptake by IGFBPs’ pathway in plaque vs control (**A**) and type A vs type B plaques (**B**). Statistical significance was calculated using robust linear modelling based on protein abundance (intensity) and detection (precursor count). Significance was accepted with an FDR-adjusted *p*-value < 0.05 (denoted with * for *p* < 0.05, ** for *p* < 0.01, and *** for *p* < 0.001).

### Culture of plaques *ex vivo* in the presence of IGF-1 results in the regulation of several proteins associated with atherosclerotic disease

Since regulation of IGF by IGFBPs was positively associated with type B plaques, we tested the effect of IGF-1 stimulation on plaques cultured *ex vivo* for 22 days, with proteomics used to quantify newly synthesized proteins (see Methods and **Fig 1**). Robust linear modelling was used to examine the differences in this parameter for plaque sections cultured in the presence versus absence of added IGF-1.

IGF-1 supplementation for 22 days, and subsequent proteomic analysis, resulted in the identification of 57 differentially synthesized proteins (52 downregulated and 5 upregulated) between the two conditions with a cut-off at a log2 fold change of ±1 (Table II). One of the top downregulated proteins in plaques treated with IGF-1 was matrix metalloproteinase 9 (MMP9; **Fig. 5A**), which is a matrix degrading enzyme that has a strong association with plaque instability and rupture [14]. MMP9 was also more abundant in type A compared to type B plaques (**Fig. 5B**). On the other hand, collagen type XXI (COL21A1) was increased in plaques treated with IGF-1.

**Figure 5.**
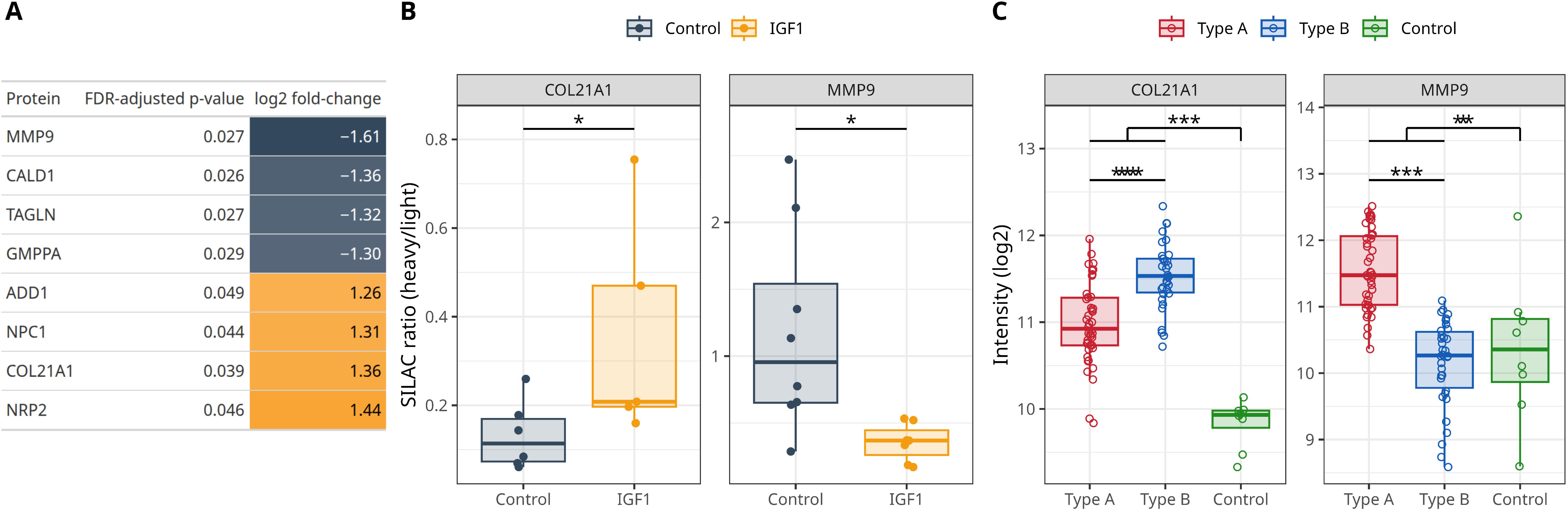
IGF-1 treatment of *ex vivo* cultured plaques (**A**) Table showing the top up-or down-regulated proteins when comparing plaques cultured with and without IGF-1. For each protein, the FDR-adjusted *p*-value and log2 fold change between the samples are shown. (**B**) Box plots displaying the SILAC ratios (heavy/light) of MMP9 and Collagen Type XXI Alpha 1 (COL21A1) in *ex vivo* cultured plaques cultured with or without IGF-1. (**C**) Box plots displaying the relative log2 intensity of MMP9 and Collagen Type XXI Alpha 1 (COL21A1) in plaque (type A and B) and control tissue. Statistical significance was calculated using robust linear modelling based on protein abundances or SILAC rations. Significance was accepted with an FDR-adjusted *p*-value < 0.05 (denoted with * for *p* < 0.05, ** for *p* < 0.01, and *** for *p* < 0.001).

## DISCUSSION

A better understanding of the disease mechanisms underlying plaque destabilisation could facilitate improved and individualised risk stratification and personalised preventive treatment strategies. The proteomic approach reported here demonstrates reproducible and consistent separation of proteins extracted from tissue samples into three clusters: control (superior thyroid) artery samples (type C), and two plaque types (types A and B). This unbiased and unsupervised proteomic analysis closely matches the macroscopic morphological assessment of the plaques, confirming the surgeons’ classification as previously reported (reference 2). As such, the type A cluster is strongly associated with soft and mixed/intermediate plaques with intra-plaque bleeding and ulceration, whereas type B is associated with the more stable morphology of ‘hard’ plaques without ulceration and haemorrhage. The precision of macroscopic classification between vulnerable and stable plaque by vascular surgeons has recently been acknowledged further and confirmed with histology [4].

These findings are consistent with our recent proof-of-concept analysis of a smaller group of plaques [2], and the data from this much larger cohort reinforce our earlier conclu-sion that the protein complement (proteomic signatures) of carotid plaques optimally partition into two groups. The group that corresponds to more stable plaques is characterized by a higher abundance of ECM structural proteins, whereas the second group primarily includes plaques with more unstable characteristics and contains elevated levels of inflammatory mediators and ECM-degrading proteases.

The inclusion of a higher number of plaques in the current study has substantially improved the data robustness reducing the potential impact of biological variability and outliers, and enhancing the statistical confidence of the clustering and differential abundance analyses. Importantly, the increased sample size allows for a better representation of intermediate (‘grey area’) plaque phenotypes, facilitating a more comprehensive characterization of plaque heterogeneity and capturing subtler molecular distinctions along the stability spectrum.

The addition of non-atherosclerotic artery samples allows a clearer differentiation of plaque-specific signatures, something that was not possible previously [2]. The current data indicate that proteins of the IGF axis are generally upregulated in plaques compared to normal vascular tissue, and that key components of the axis (IGF-1, IGF-2, IGFBP-1 and IGFBP-2) were only present in plaques. This is supported by previous studies investigating IGF-1, IGF-1R and IGFBP-1-5 levels in coronary atherectomy specimens, where IGF-axis proteins could be detected in plaques but not in non-atherosclerotic coronary arteries [15].

Another recent study of human carotid atherosclerotic plaques [16] also identified proteins related to the IGF-axis (IGF-2, IGF-2R, IGFALS, and IGFBP-3, 5, 6 and 7) with these being generally more abundant in the plaque core compared to its periphery, and with IGF-2, IGFBP-3 and IGFBP-5 more abundant in calcified compared to non-calcified plaques. However, this study utilized a stepwise extraction protocol, and the data varied between different fractions [15]. Together, these data support the hypothesis that IGF-axis proteins are abundant in carotid (and coronary) plaques when compared to non-plaque tissue, and that IGF-axis proteins are more abundant in stable/calcified plaques in general. These results also support the assertion that IGF signalling may enhance plaque stability in advanced atherosclerotic disease [17].

Circulating levels of IGF axis proteins have been previously implicated in cardiovascular disease, and serum IGF-1 levels have been investigated as a potential predictor of risk. Low serum IGF-1 have been associated with an increased risk of ischemic heart disease and stroke, with high serum IGF-1 levels being protective [18–20]. While increased levels of IGF axis proteins in plaques are likely to at least partially stem from influx from the circulation, there is also evidence for increased *in situ* synthesis of these proteins within the plaques as mRNA for both IGF-1 and –2, as well as IGFBPs have been detected in both coronary [21] and carotid [15] atherectomy specimens.

Binding of IGF-1 (or IGF-2) to IGF-1R stimulates intracellular signal transduction, which induces cell survival and growth [17]. On the other hand, IGF-2R is a scavenger receptor that mediates IGF internalization and degradation [22]. We detected significantly more IGF-1R, and significantly less IGF-2R in plaques with stable morphology (type B) compared to those associated with unstable morphology (type A). This could indicate that the level of IGF signalling is a differentiating factor between stable and unstable plaques, as supported by animal studies, where enhanced IGF-1 levels induced features associated with plaque stability, including increased fibrous cap area, SMC and collagen content, and a decrease in plaque burden [13],[23–25]. Both SMC- and macrophage-specific IGF-1 overexpression (individually) in *ApoE^-/-^* mice has been shown to increase features of plaque stability [26, 27], whereas macrophage-specific IGF-1R deficiency in *ApoE^-/-^* mice accelerated atherosclerosis and induced plaques with an unstable morphology [28]. All three models provided evidence for increased plaque collagen content in the presence of (increased) IGF-1 signalling. This is in line with our data showing downregulation of MMP9 and upregulation of COL21A1 in response to IGF-1 treatment in *ex vivo* cultured plaques. MMP9 is a collagen-degrading protease and its activity is associated with plaque rupture [29]. Using spatial transcriptomics Sun et al. have shown that MMP9 expression is increased near the rupture site of carotid artery plaques and that MMP9 expression is higher in plaques from symptomatic patients [30]. Additionally, pigs with familial hypercholesteremia, supplemented with IGF-1, exhibited lower expression of MMP9 at the mRNA level in addition to a more stable plaque phenotype [25]. Decreased MMP9 transcription and protein expression could result in decreased MMP9 activity, which would result in decreased collagen degradation and increased fibrous cap area and collagen content in plaques. Thus, decreased MMP9 provides a mechanistic link between IGF-1 and plaque stabilization.

In our *ex vivo* model we retain the complexity of the human atherosclerotic plaque including different cell types, lipids and extracellular matrix. *Ex vivo* culture separates the plaque from the circulation, which allows us to focus on intraplaque mechanisms, and we show that these can be altered through supplementation with IGF-1. Our model brings the testing of IGF-1 as a plaque-stabilizing therapy one step closer to patients, while demonstrating the promise of applying *ex vivo* culture in pre-clinical drug testing. In the current study, we identified a limited number (57) of differentially synthesised proteins, which is likely due to the high variability of plaque subtypes, ranging from unstable to stable with intermediate plaque morphologies. Classification of plaques before *ex vivo* culture in addition to an increased sample size would probably enhance the observed effects, since our data indicate that IGF-1 effects are plaque subtype dependent.

In conclusion, this study provides compelling evidence for an involvement of the IGF axis in carotid atherosclerosis and plaque stability. Our data, together with previous reports of the plaque stabilising effect of IGF-1, indicates that drugs targeting the IGF axis might be appropriate to prevent acute cardiovascular events.

## Acknowledgements

This work was supported by grants from Novo Nordisk Foundation (NNF20SA0064214 to M.J. Davies; NNF22OC0079176 to L.G. Lorentzen) and the Novo Nordisk Foundation-University of Copenhagen BRIDGE scheme (to L.G. Lorentzen).

