## Supplemental material for "Proteomics and Ex Vivo Plaque Culture Identify the Insulin-Like Growth Factor Axis as a Regulator of Carotid Plaque Stability"

**Supplementary data**

**SUPPLEMENTARY METHODS**

- Arg10 / Lys8: Heavy isotope-labeled arginine (+10 Da) and lysine (+8 Da)
- CAA: 2-chloroacetamide
- DDA: Data-dependent acquisition
- DDA-PASEF: Data-dependent acquisition with parallel accumulation–serial fragmentation
- DIA: Data-independent acquisition
- DIA-NN: Data-Independent Acquisition by Neural Networks (software)
- DIA-PASEF: Data-independent acquisition with PASEF
- FBS: Fetal bovine serum
- FDR: False discovery rate
- IGF-1: Insulin-like growth factor 1
- Lys-C: Endoproteinase Lys-C
- m/z: Mass-to-charge ratio
- MS: Mass spectrometry
- MS/MS: Tandem mass spectrometry
- PBS: Phosphate-buffered saline
- PASEF: Parallel accumulation–serial fragmentation
- PSM: Peptide–spectrum match
- SILAC: Stable isotope labeling with amino acids in cell culture
- TCEP: Tris(2-carboxyethyl)phosphine
- TFA: Trifluoroacetic acid

***Ex vivo* plaque culture**

The *ex vivo* plaque culture method was developed based on that described by Lebedeva et al.^1^. Carotid artery plaques removed by endarterectomy were placed in RPMI 1640 medium (Gibco, Thermo Fisher Scientific, Waltham, MA, USA, cat. 12633012), containing 10% FBS (Gibco, Thermo Fisher Scientific, cat. 10270106), 2 mM GlutaMAX (Gibco, Thermo Fisher Scientific, cat. 35050061), 1x penicillin-streptomycin (10,000 Units/mL; Gibco, Thermo Fisher Scientific, cat. 15140122) and 1x amphotericin B (Gibco, Thermo Fisher Scientific, cat. 15290026) and were transported to the laboratory in a portable incubator (Cellbox Ground 2.0, Cellbox Solutions GmbH, Hamburg, Germany) set to 37 ^°^C and 5% CO_2_. Plaques were cut into cross-sectional rings (~2-3 mm) and washed in sterile PBS, before transfer to wells containing SILAC medium (Thermo Fisher Scientific, cat. A33973) prepared according to manufacturer’s protocol. Plaques were cultured under 37 ^°^C and 5% CO_2_ for 22 days, with the medium being changed on day 1 followed by every third day. Supplementation with 100 ng/ml IGF-1 (Bio-techne (R&D Systems), Minneapolis, MN, USA, cat. 291-G1-200) was started on day 0, and new IGF-1 was added every time the medium was changed.

**Sample processing and spectral library generation**

Protein extract from 6 plaques was digested as described by Doellinger et al. ^2^. Briefly, proteins were reduced and alkylated with TCEP and CAA at 95°C for 10 min, before Lys-C (1:100) and trypsin (1:50) was added to the protein extracts and incubated overnight at 37°C. The resulting peptides were delipidated with tert-methyl butyl ether before being fractionated by high-pH reverse phase chromatography. The peptide species (50 μg) were loaded onto a Kinetex EVO C18 column (1×150 mm, 1.7um particle size, Phenomenex) size column, and separated using a 50 min binary gradient (5% mobile phase B for 5 minutes followed by an increase to 40% mobile phase B in 45 minutes; mobile phase A: 20 mM ammonium formate, pH 10.5; mobile phase B: 80% acetonitrile with 20 mM ammonium formate, pH 10.5). Eluates were collected into Eppendorf LoBind plates every minute using an Opentron OT2 robot. Eluates from 0-12 min were pooled with eluates from 36-48 min, resulting in a total of 36 fractions per sample. Fractionated peptides were then dried down by vacuum centrifugation before being resuspended in 0.1% TFA in MS water and separated using a Dionex Ultimate RSLCnano system (Thermo Scientific) equipped with an Aurora C18 column (25 cm×75 μm, 1.6 μm particle size, IonOpticks), maintained at 50 °C and coupled to a timsTOF Pro mass spectrometer (Bruker Daltonics, Bremen, Germany). Peptides were eluted using a 100 min binary gradient (2-35% mobile phase B; mobile phase A: 0.1% v/v formic acid in MS water; mobile phase B: 0.1% v/v formic acid in acetonitrile) with data acquisition in DDA-PASEF mode^3^.

We combined all 216 DDA runs to generate a spectral library using the FragPipe software suit (v. 22.0)^4^. The MSFragger algorithm (v. 4.1)^5^ was used to search MS/MS spectra against the SwissProt database of human protein sequences, supplemented with common contaminant protein sequences (downloaded 06-12-24) and reversed sequences as decoys. Cys alkylation (*m/z* +57.02) was set as a fixed modification, and Met oxidation (*m/z* +15.99) and protein N-term acetylation (*m/z* +42.01) were set as variable modifications. Tryptic peptides with a maximum of two missed cleavages and length between 7 and 50 amino acids were allowed. MSBooster (v. 1.2.31)^6^ was used to rescore peptide-spectrum matches (PSMs) before validation of the PSMs with Percolator (v. 3.6.5). The Philosopher software suit (v. 5.1.1)^7^ was used for protein inference (ProteinProphet) and filtering of the search results to obtain a 1% false discovery rate at protein and peptide levels. Finally, a spectral library was generated from the remaining PSMs using EasyPQP.

For analysis of SILAC labelled samples (*ex vivo* cultured plaques), a predicted library was constructed from the DDA library with Arg10/Lys8 labels included using DIA-NN (v. 2.0)^8^.

These libraries were then used in DIA-NN to analyse DIA-PASEF files. DIA-NN was configured to allow peptides with 7-35 amino acids, and precursors between *m/z* 300 - 1100 (charge state 2-4). A single missed cleavage was allowed, and Cys alkylation was set as a fixed modification. Quantification was performed using the QuantUMS algorithm^9^ and with match-between-runs enabled.

In the case of SILAC labelled samples, DIA-NN was run with these additional parameters to allow for SILAC quantification: --fixed-mod SILAC,0.0,KR, label --lib-fixed-mod SILAC --channels SILAC,L,KR,0:0;SILAC,H,KR,8.014199:10.008269 --original-mods --channel-run-norm
